# Interruption of rat absence seizures by auditory stimulation

**DOI:** 10.1101/2025.10.10.681637

**Authors:** Cian McCafferty, Xinyuan Zheng, Renee Tung, Benjamin F. Gruenbaum, Hal Blumenfeld

## Abstract

Absence seizures are episodes of impaired consciousness and responsiveness that impact an individual’s ability to interact with the world around them. Childhood absence epilepsy, a condition defined by these seizures, can have profound effects on children’s social, educational, and psychological development. Absence seizures are accompanied by a distinctive electrographic signature called a spike-wave discharge. The impairment of consciousness associated with a spike-wave discharge can be variable: some people maintain responsiveness during some absence seizures, and some rodent oscillations resembling spike-wave discharges may not have any behavioral impact. We previously observed that spike-wave discharges in the Genetic Absence Epilepsy Rat from Strasbourg model sometimes terminated shortly after presentation of a conditioned auditory stimulus. In this study we found that these terminations were caused by the stimuli and that they occurred after approximately 50% of stimuli. We also found that the probability of a spike-wave discharge being interrupted depended on stimulus timing, degree of conditioning, and electrographic signal power. These data provide insight into the factors that determine the mechanisms of absence seizure termination, with possible implications for therapy.

## Introduction

Absence seizures are episodes of impaired consciousness accompanied by a distinctive spike-and-wave discharge (SWD) signature on electroencephalography. They tend to last tens of seconds at most and are the defining seizure type of childhood (CAE) and juvenile (JAE) absence epilepsies, as well as a commonly observed seizure type in other epilepsy syndromes and presentations (Fisher et al., 2017). The impaired consciousness during an absence seizure constitutes an interruption in the person with epilepsy’s life – it abolishes or severely limits responsiveness, learning, and development for its duration (Blumenfeld, 2005a). Children with CAE or JAE can have hundreds of such episodes daily, and these syndromes are associated with developmental, education, and psychiatric challenges (Shinnar et al., 2017; Gruenbaum et al., 2021).

The nature of the impairment of consciousness during an absence seizure is not fully established. While clinical descriptions of absence seizures tend to suggest a complete loss of consciousness, there is longstanding if incomplete evidence from human and animal studies that SWDs do not always fully abolish responsiveness (Mirsky and Vanburen, 1965; Rajna and Lona, 1989). Children with CAE can sometimes continue carrying out a pre-set task during a seizure and even respond selectively to stimulation (Guo et al., 2016). Rodents exhibiting SWDs, or similar electrographic oscillations, have been shown to experience shorter SWDs when each one was followed with a food reward (Taylor et al., 2017). Oscillations resembling SWDs have also been observed and proposed as physiological states in multiple strains of inbred, outbred, and even wild rats (Wiest and Nicolelis, 2003; Shaw, 2004; Taylor et al., 2019). Although both thalamus and cerebral cortex are heavily involved in SWDs, expressing strong rhythmic activity on aggregate, it has been shown that at least some information processing from sensory stimulation is preserved during seizures (Chipaux et al., 2013; Williams et al., 2016) and that not all neurons in these areas express rhythmic patterns of firing during seizure (McCafferty et al., 2023). Understanding the nature and mechanisms of impairment of consciousness during absence seizures may allow better care of children with CAE including titration of anti-seizure medications (Kessler and McGinnis, 2019) and point towards improved treatments to address the lack of progress in this area since the adoption of ethosuximide as the gold-standard treatment (Brigo et al., 2021).

In summary, impairment of responsiveness and consciousness during absence seizures is variable. Previous studies have begun to describe this variability (Berman et al., 2010; Taylor et al., 2017), but we do not know what underlies it – i.e. what causes one absence seizure to fully impair consciousness while another allows continued responsiveness. Guo et al. found that seizures impairing behavioral responses had larger associated electroencephalographic and fMRI changes (Guo et al., 2016), and we wanted to investigate whether a similar relationship existed in an established rodent model of absence seizures to facilitate future cellular and subcellular mechanistic investigations.

In particular, the present study focuses on the variable ability to *interrupt* an absence seizure with external sensory stimulation, as a way to assess the variable behavioral severity of seizures. Interruption of human absence seizures by auditory or somatosensory stimulation has been described in several clinical reports (Jung, 1962), including a study that found the interruption of absence seizures by auditory stimuli was more effective if the stimulus occurred at earlier times (first 3 seconds) during the SWD (Rajna and Lona, 1989). Vigorous tactile stimulation has been reported to interrupt tonic-clonic seizures in children (Pothman, 1982). Increased vigilance level and engagement in behavioral tasks has long been known to suppress the occurrence of generalized spike-wave discharges and absence seizures in human patients (Lennox et al., 1936; Gastaut, 1954; Kooi and Hovey, 1957; Tizard and Margerison, 1963; Davidoff and Johnson, 1964; Bureau et al., 1968; Sellden, 1971; Horita et al., 1991; Opp et al., 1992; Berman et al., 2010; Zarowski et al., 2011) as well as in animal models (Coenen et al., 1991; Van Luijtelaar et al., 1991; Vergnes et al., 1991; Smyk et al., 2011). Caregivers in another study reported that some seizures can be interrupted by physical stimulation although the study did not specify the seizure type (Pinikahana and Dono, 2009). Other studies report that patients can control or prevent their own seizures by cognitive or physical actions, again without specifying seizure type (Spector et al., 2000; Kotwas et al., 2016; Leeman-Markowski and Schachter, 2017). Olfactory stimulation can sometimes interrupt temporal lobe seizures (Gowers, 1881; Efron, 1956, 1957; Betts, 2003). Focal interictal epileptiform spikes can be suppressed by somatosensory stimuli or motor activity (Tassinari, 1968; Ricci et al., 1972). In summary, it has long been known that external sensory stimuli may modulate human generalized or focal epileptic activity, possibly through a modulation of arousal state, but the mechanisms of this modulation have not been investigated in detail.

Understanding absence seizure interruption would be greatly facilitated by investigation in experimental animal models. Previous studies have shown that auditory or other sensory stimuli can interrupt SWDs in rat models (Vergnes et al., 1982; Drinkenburg et al., 2003; Wiest and Nicolelis, 2003; Shaw, 2004; Pearce et al., 2014; Rodgers et al., 2015; Taylor et al., 2017). But factors that are related to susceptibility of absence seizures to interruption have not been studied in detail. We also wished to provide further answers to the outstanding question of whether rodent spike-wave discharges are a useful model of absence seizures (Shaw, 2004; Chipaux et al., 2013; Stenroos et al., 2024) or rather reflect a physiological process specific to the rodent brain with limited applicability to epilepsy (Wiest and Nicolelis, 2003; Taylor et al., 2017; Taylor et al., 2019; Taylor et al., 2022).

We therefore made use of a dataset we previously collected in an investigation of cortical and thalamic neuronal activity during rat absence seizures, in which a classical auditory conditioned response task was modified by decreasing stimulus intensity (to 45 dB) and increasing average inter-stimulus interval until rats expressed seizures during recording sessions (McCafferty et al., 2023). Rats were trained to associate the auditory stimulus with a sucrose-water reward, and stimuli were then presented both between and during seizures. This analysis concerns the latter set of stimuli – those delivered during an absence seizure. We had anecdotally observed that some absence seizures terminated shortly following these stimuli and wished to investigate whether there was a causal relationship between stimulation and termination.

We found that approximately half of all stimuli delivered during a seizure resulted in a premature termination of that seizure – within one second of stimulus presentation. Behavior, as indicated by lick rate at the reward spout, recovered within ten seconds of this interruption. Interruptions were more likely to occur when a stimulus was delivered earlier in a seizure, suggesting the processing and effects of incoming sensory information may vary during the seizure. The proportion of interrupted seizures in a recording session positively correlated with the number of previous sessions a rat had undergone, suggesting the salience of the stimulus or the degree of task learning can influence responsiveness during seizure. Interrupted seizures had different electrographic properties to those that were not interrupted, but we did not find any long-timescale pre-seizure indicators of whether a seizure might be interrupted or not.

These results indicate some influencing factors and characteristics of variability in absence seizure impairments of consciousness or responsiveness. The observations are similar to those in people with absence seizures, in that the events are capable of impairing a rewarding and appropriate behavior, but are not a monolithic abolition of responsiveness. Future studies might explore the neuron- and network-level mechanisms determining this variability and the nature of the variability in people with absence seizures, as well as in other animal models.

## Methods

Experiments were carried out under approval by the Yale University Office of Animal Research Support. Genetic Absence Epilepsy Rats from Strasbourg (GAERS) between 3 and 7 months of age were used. Animal husbandry, surgical implants, and behavioral paradigms were previously described in (McCafferty et al., 2023). Female animals were used because of requirements specific to the original study (McCafferty et al., 2023). They had access to food and water ad libitum except where noted below, and were on a 12:12h light:dark cycle. Animals were group housed prior to surgery and single-housed thereafter to prevent injury or damage to implants.

### Surgeries

Electroencephalography electrodes were implanted under isoflurane anesthesia, with induction at 5% isoflurane in oxygen at 2 liters/minute flow rate in a plexiglass chamber and maintenance at 1.5-2% at 1 liter/minute flow via a face mask and scavenging suction. Surgical plane of anesthesia was verified by monitoring of breathing rate and pedal reflex. Analgesia was provided by two intraperitoneal carprofen injections at 5 mg/kg the first at 1h before surgery and the second at 24 h post-surgery in order to provide 48 hours of pain relief.

Bilateral frontal, parietal and cerebellar epidural screw electrodes (0-80 x 3/32, PlasticsOne) were implanted with a plastic pedestal connector (PlasticsOne). Fronto-parietal differential EEG was collected in each hemisphere from these electrodes via a Model 1800 Microelectrode AC Amplifier & Headstage (A-M Systems) between 0.1 and 1000 Hz with 1000x gain, using the cerebellar electrode as ground. Signals were digitized at 1000 samples/second with a Micro1401 acquisition unit (Cambridge Electronic Design) and saved in Spike2 v8 (Cambridge Electronic Design). Transistor-transistor logic (TTL) pulses from behavioral Med-PC V Software (Med Associates, Inc.) were send to Spike2 to allow alignment of behavioral and EEG data.

### Behavioral Training

The behavioral paradigm employed here was described in (McCafferty et al., 2023) as a “sensory tone detection task” and is summarized here. Animals were switched from ad libitum to restricted food access until they reached 90% of their initial body mass in order to increase their motivation to acquire sucrose water rewards, due to their decreased baseline levels of such motivation relative to non-epileptic rats (Jones et al., 2008). Restriction was terminated when rats completed training stage 4 (below) and at least 1 week of ad libitum access was allowed prior to surgery.

Training consisted of 5 stages, from 0 to 4. Stage 0 involved introduction of 20% sucrose (weight/volume) in water, manually dispensed from a 1 ml syringe (Becton Dickinson) in the animals’ home cage in 10-minute sessions repeated up to 4 times daily. This stage ended when rats consumed 2 ml of water within a session. Stage 1 was the same as Stage 0 but took place in a custom operant chamber (Med Associates Inc) instead of the home cage and consisted of a single session. All subsequent stages of training and testing took place in the custom operant chamber (Figure 1A). In stage 2 the sucrose water (90 μL) was instead provided by a delivery port in the operant chamber and delivered at the same time as a 0.5 second or 1.0 second 8 kHz 80 dB tone. Because initial analyses showed very similar results for 0.5 and 1.0 second tones, results for both stimulus durations were subsequently combined in all analyses. The sucrose water delivery occurred at 60 s intervals with no action required by the animal. In stage 3 the sucrose water delivery required the rat to lick at the spout within 10 seconds of tone presentation. In Stage 4 the intensity of the stimulus was decreased to 45 dB and the inter-stimulus interval was increased to a random range from 150 to 120 s, or the automated detection of a SWD (McCafferty et al., 2023), whichever was sooner.

**Figure 1:**
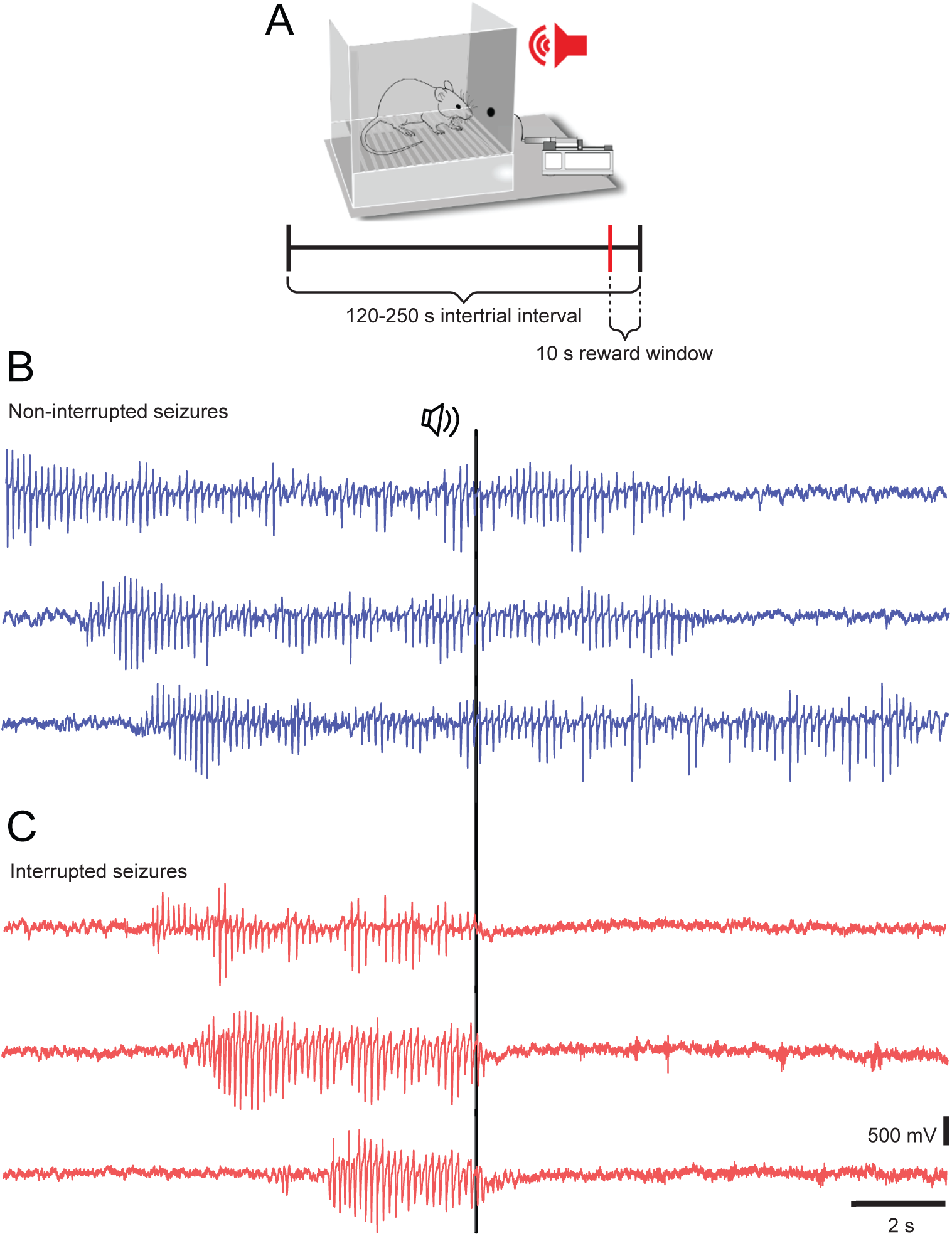
Illustration of experimental setup and EEG of non-interrupted and interrupted seizures. A. Schematic of the experimental setup. B. Examples of EEG recordings from seizures that did not end soon after the presentation of the auditory stimulus. C. Examples of seizures that ended within 1 second after the stimulus.

Each training stage from 2 to 4 featured sessions up to 2 hours duration and was considered complete when a rat received 50 rewards within one session. Training from stage 0 to 4 typically took 2 weeks to complete.

### Behavioral Testing

After completion of training, a 1-week rest period, EEG implantation, and a further 5-day recovery, animals had a single refamiliarization session at stage 4. Testing then began at day 7 post-surgery with 1 daily session per animal. The testing protocol was identical to stage 4 training. The auditory tone was now considered a conditioned stimulus. Stimuli were delivered during seizures and during inter-seizure periods. For this study we analyzed stimuli delivered during seizures only.

### Spike-wave discharge labelling and EEG processing

Spike-wave discharges were labeled using Spike2 software as previously described (McCafferty et al., 2018). Smooth (moving average) and DC removal (subtraction of slow changes) functions were used to remove high- and low-frequency portions of the signal that did not overlap with the spike-wave range. Spike-wave discharges were then labeled semi-automatically using a combination of expected amplitude, frequency, and temporal characteristics as previously described (McCafferty et al., 2018).

### Spectral analyses

When the power in certain frequency bands was of interest, EEG signals were first prepared by user identification and removal of artifact periods in Spike2. To visualize power changes around seizure start (t ± 5 s, with t being the seizure start time, see Figure 3C), seizure end (t ± 5 s, with t being the seizure end time) (Fig. 3C) and stimulus onset (t ± 2 s, with t being the stimulus onset time) (Fig. 3D), the root-mean-square voltage (VRMS) was calculated in consecutive overlapping 1 s time bins at the original sampling rate (1000 Hz) for each seizure. To analyze the time course changes in VRMS, the baseline period was defined as from 5 to 1 s before the initiation of each individual seizure and mean percent changes of VRMS values were computed by normalizing VRMS values using [(VRMS signal – mean VRMS baseline period)/mean VRMS baseline period] X 100%. Spectrograms were also derived from the EEG signals using MATLAB’s spectrogram function, with a window size of 250 ms and a sampling rate of 1000 Hz. Analyses of long-term state changes in power prior to seizures were conducted by calculating the mean power from 120 to 1 s before the initiation of each individual seizure and then normalizing power values by dividing them by the mean over the same time range. The resulting values were log-transformed to express power in dB. The baseline-corrected, log-transformed spectrograms were averaged across seizures within an animal before being averaged across animals. The time course of power in frequency bands of interest was calculated similarly, except that values were averaged over the relevant frequency ranges.

### Statistical analyses

Data are described using mean across seizures and standard error of the mean across seizures except if explicitly indicated. Each seizure was considered a sample (n = 427 seizures: n*_interrupted_* = 200 seizures, n*_non-interrupted_* = 227 seizures). Differences between the interrupted and non-interrupted groups were tested by using two-sample t-tests. The Kolmogorov-Smirnov test was used to determine that the distribution of stimulus or seizure counts over time is non-uniform. To model the potential correlation between the number of training sessions an animal received and the proportion of interrupted seizures in that session, we used a linear regression to fit the data through the least squared method and reported the resulting Pearson’s R and associated p-value.

## Results

### Establishing interruption of absence seizures

While investigating the ability of GAERS to respond to auditory conditioned stimuli (Fig. 1A) during SWDs, we noticed that some of those SWDs terminated within a few seconds of stimulus presentation (Fig. 1B,C). We therefore hypothesized that the stimuli were causing the premature termination of the SWD: interrupting the seizure. To investigate whether this was indeed the case, and to distinguish seizure interruption from spontaneous or coincidental termination, we investigated the distribution of latencies between stimulus presentation and seizure termination. We reasoned that, if stimulus presentation and seizure termination were unrelated, the distribution of times from stimulus presentation to seizure end across seizures should be relatively uniform. For this investigation we selected those SWDs that had a stimulus delivered within their duration, discounting SWDs without stimuli and stimuli that were delivered outside SWDs. Henceforth the terms “seizure” and “SWD” will be used interchangeably and refer to this subgroup of all events recorded.

Contrary to this null hypothesis, we found that approximately half of all seizures (200) had a stimulus 1 second or less before their termination (Fig. 2A). The other half of seizures (227) had stimuli distributed from 40 seconds to 2 seconds before termination, with a skew towards shorter latencies (mean latency 4.1 +/- 6.9 s). It was possible that this skew, and the concentration of stimuli in the second before termination, was due at least in part to the distribution of seizure durations: seizures averaged 8.5 +/- 10.1 seconds in duration (Fig. 2B) with a similar skew towards shorter durations. To account for this possibility, we divided the number of stimuli falling in each time bin relative to termination by the number of seizures of at least that duration. A conservative estimate of the number of seizures expected to end spontaneously within a certain time period was obtained by adding up the total number of seizures up to that duration. The number of stimuli falling between -1 and 0 seconds before termination was divided by the number of seizures lasting at least 1 second in duration (i.e. all seizures); the number of stimuli between -2 and -1 seconds before termination was divided by the number of seizures lasting at least 2 seconds, and so on. The resulting distribution (Fig. 2C) was not uniform (p = 4.3 e-7, K-S test), as would have been expected if stimulus time and seizure termination were unrelated. Instead, a large peak was apparent in the 0 to 1 second bin.

**Figure 2:**
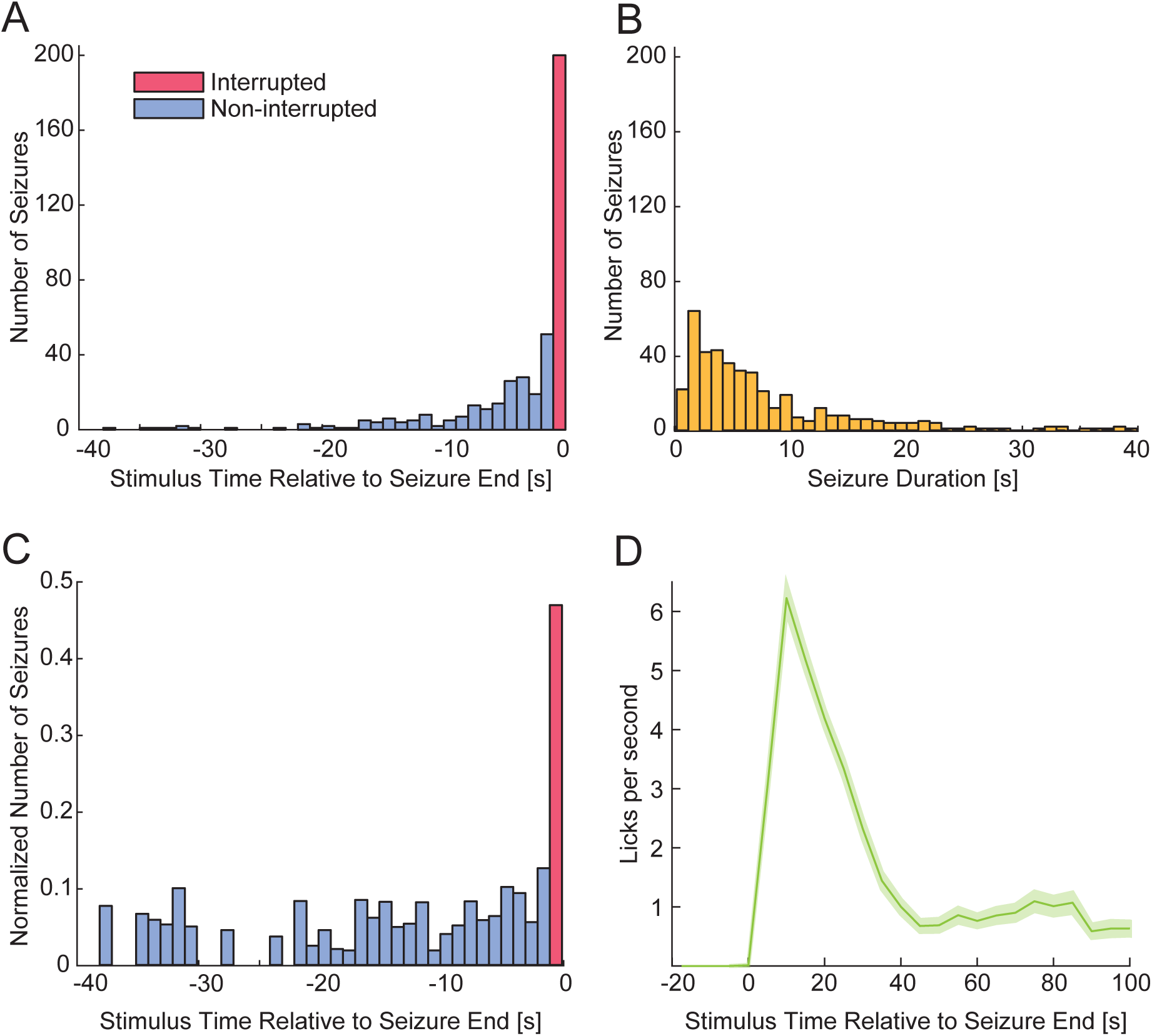
Conditioned auditory stimuli interrupt approximately 50% of seizures electrographically and behaviourally. A: Distribution of stimulus time to seizure end. Mean: 4.1 +/- 6.9 s. Interrupted seizures (red) are defined as those terminating within 1s of an auditory stimulus onset. B: Distribution of seizure durations. Mean: 8.5 +/- 10.1 s. C: Frequency distributions of stimulus timing relative to the end of a seizure (A), and of seizure durations (B), are combined to obtain the normalized probability of a seizure falling in each time bin relative to the seizure end (see Methods for details). Kolmogorov-Smirnov test indicates that the distribution of stimulus count/seizure count over time is not uniform (p = 4.3 e^-7^). D: Behavioral licking response rate mean time course (± SEM) across interrupted seizures aligned to time of seizure end.

Based on these findings we labelled those seizures terminating within 1s of the stimulus as “interrupted”, and their counterparts “non-interrupted”. While the interrupted seizures were defined based on the cessation of the electroencephalographic SWD, we were interested in whether the rat’s behavior, as well as its electroencephalography, was restored upon interruption. The auditory stimulus heralded the availability of a sucrose-water reward at a spout in the operant chamber (Fig. 1A), and rats were motivated to lick at that spout when possible due to their training.

We therefore quantified the lick rate per second after interrupted seizures and found a recovery to approximately 6 licks per second within 10 seconds of termination, indicating that rats were able to respond following interrupted seizures (Fig. 2D) similarly to their ability at non-seizure baseline periods (e.g. see McCafferty et al., 2023, Fig. 2C).

### Factors influencing interruptions

We next investigated factors that could explain whether a stimulus would interrupt a seizure, starting with factors extrinsic to the seizure itself: the timing of the stimulus and the training state of the animal. We found that, from very similar samples (200 each), more interrupted seizures than non-interrupted seizures had stimuli falling in the first second of the seizure (red bar visible), while more non-interrupted seizures had stimuli falling later in the seizure (blue bars visible) (Fig. 3A). This is reflected in a lower mean time from seizure onset to stimulus onset in the interrupted group. The mean latency for interrupting stimuli was 3.5 +/- 4 s and for non-interrupting was 5.2 +/- 6.8 s (t-test comparing latencies p = 0.0023). We considered that the salience of the stimulus to the animal might influence its ability to “penetrate” a seizure state (Chipaux et al., 2013; Stenroos et al., 2024), and that this salience might increase with repetition of the behavioral paradigm. We found that there was a positive correlation (r = 0.506, p = 0.027) between the number of sessions an animal had experienced and the proportion of seizures in that session that were interrupted, averaged across animals (Fig. 3B). These results show that the propensity of a seizure to be interrupted by a stimulus can be influenced by the timing of that stimulus and, potentially, the altered meaning of the stimulus for the animal during training.

Factors intrinsic to a seizure may also plausibly have influenced the ability of a stimulus to cause its premature termination. We therefore analyzed the VRMS magnitude of the EEG signals for interrupted and non-interrupted seizures. Considered over the entirety of the seizure, the mean percentage change (vs. baseline) in VRMS of the spike-wave discharge was higher in non-interrupted seizures (Interrupted: 224.05% +/- 9.76%; Non-interrupted: 260.77% +/- 9.42%; p = 0.0073) (Fig. 3C). We also considered that the immediate temporal environment of the stimulus could influence its effects, as seizures wax and wane in intensity over their duration. In the 250 ms immediately preceding stimuli, the mean VRMS change was higher in non-interrupted seizures (Interrupted 224.76% +/- 9.97%; Non-interrupted 269.58% +/- 9.95%; p = 0.0016) (Fig. 3D). These results suggest that the electrophysiological features of a seizure, both overall and immediately, may influence the effect of the seizure on an auditory stimulus. The decreased EEG power of interrupted seizures may therefore indicate that their lower physiological severity.

**Figure 3:**
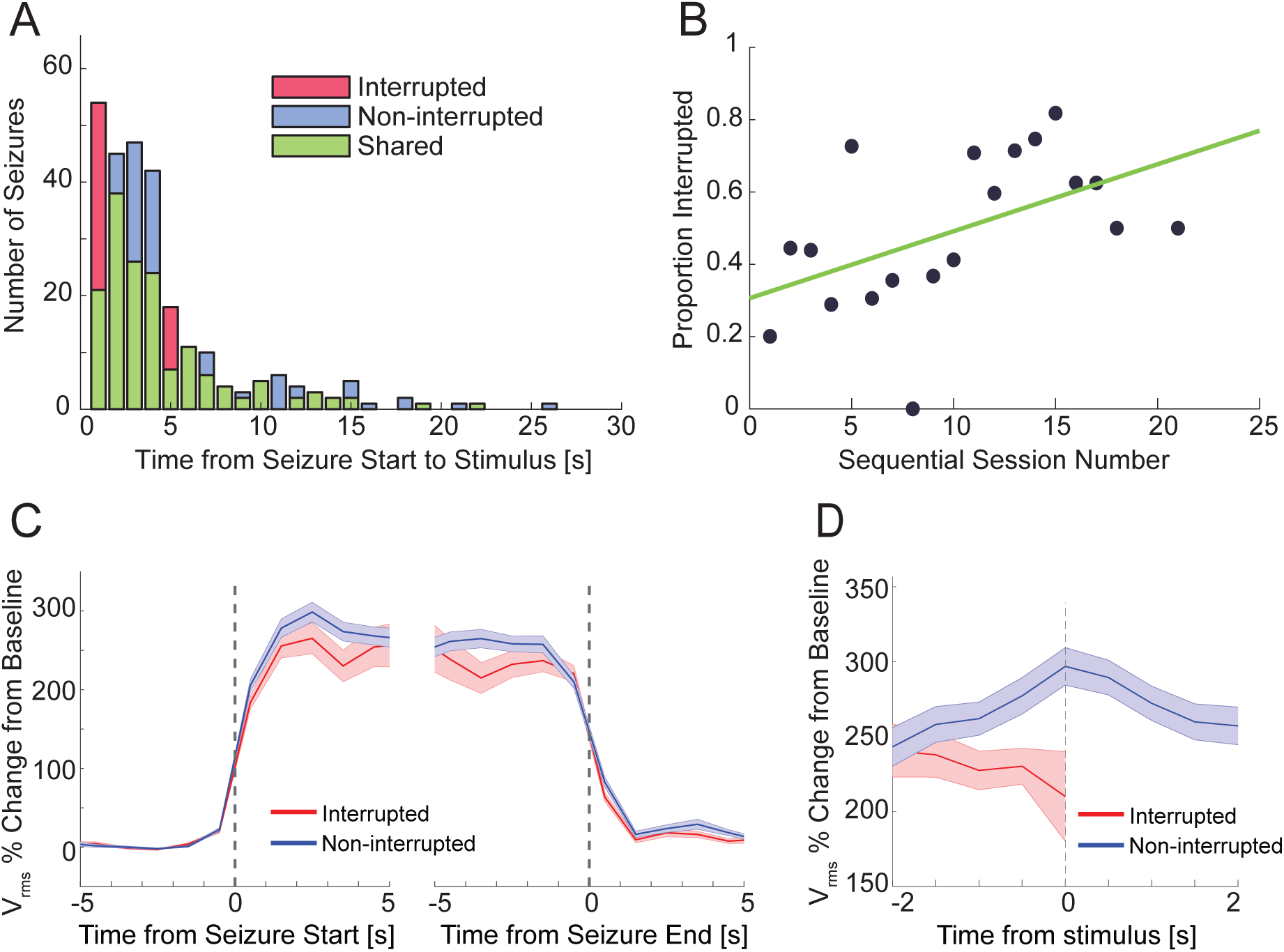
Seizures are more likely to be interrupted by earlier stimuli, in trained animals, and when VRMS power is lower. A: Distribution of latencies between the start of a seizure and an auditory stimulus delivered within that seizure shows that “interrupting” stimuli were more likely to fall earlier in a seizure than were “non-interrupting” stimuli. The mean latency for interrupting stimuli was 3.5 +/- 4 s and for non-interrupting was 5.2 +/- 6.8 s (t-test comparing latencies p = 0.0023). B: There was a positive correlation between the number of behavioral testing sessions (after training completion) an animal had experienced and the proportion of seizures in that session that were interrupted (r = 0.506, p = 0.027). C: EEG signals from interrupted seizures exhibited different power characteristics compared to non-interrupted seizures. The mean VRMS percentage change during ictal period was higher for non-interrupted seizures (t-test comparing VRMS changes p = 0.0073). D: The mean VRMS change just prior to the stimulus was higher in non-interrupted seizures (t-test comparing VRMS changes p = 0.0016).

The final set of factors we considered as potential influencers of seizure interruption related to the animal’s general state prior to the seizure. While absence seizures can arise from various initial arousal states, their properties vary according to state (Sadleir et al., 2008), and we proposed that seizures originating in different states may not be equal in terms of their susceptibility to interruption. However, we did not find a difference in the power spectra of the EEG signals in the 120 seconds preceding interrupted and non-interrupted seizures (Fig. 4A), nor did we find a difference in any ten-second time bins from the time courses of power in low (1 – 39 Hz) and high (40 – 500 Hz) bands (Fig. 4B,C). As the animal’s arousal state may also be reflected in its behavior, we also quantified time courses of lick rate in the same window and again did not find a difference between our groups (data not shown). These results suggest that, if any determinants of whether a seizure will be interrupted are present before its initiation, they are not reflected in the fronto-parietal EEG signals or lick rate data that we had available before seizure onset.

**Figure 4:**
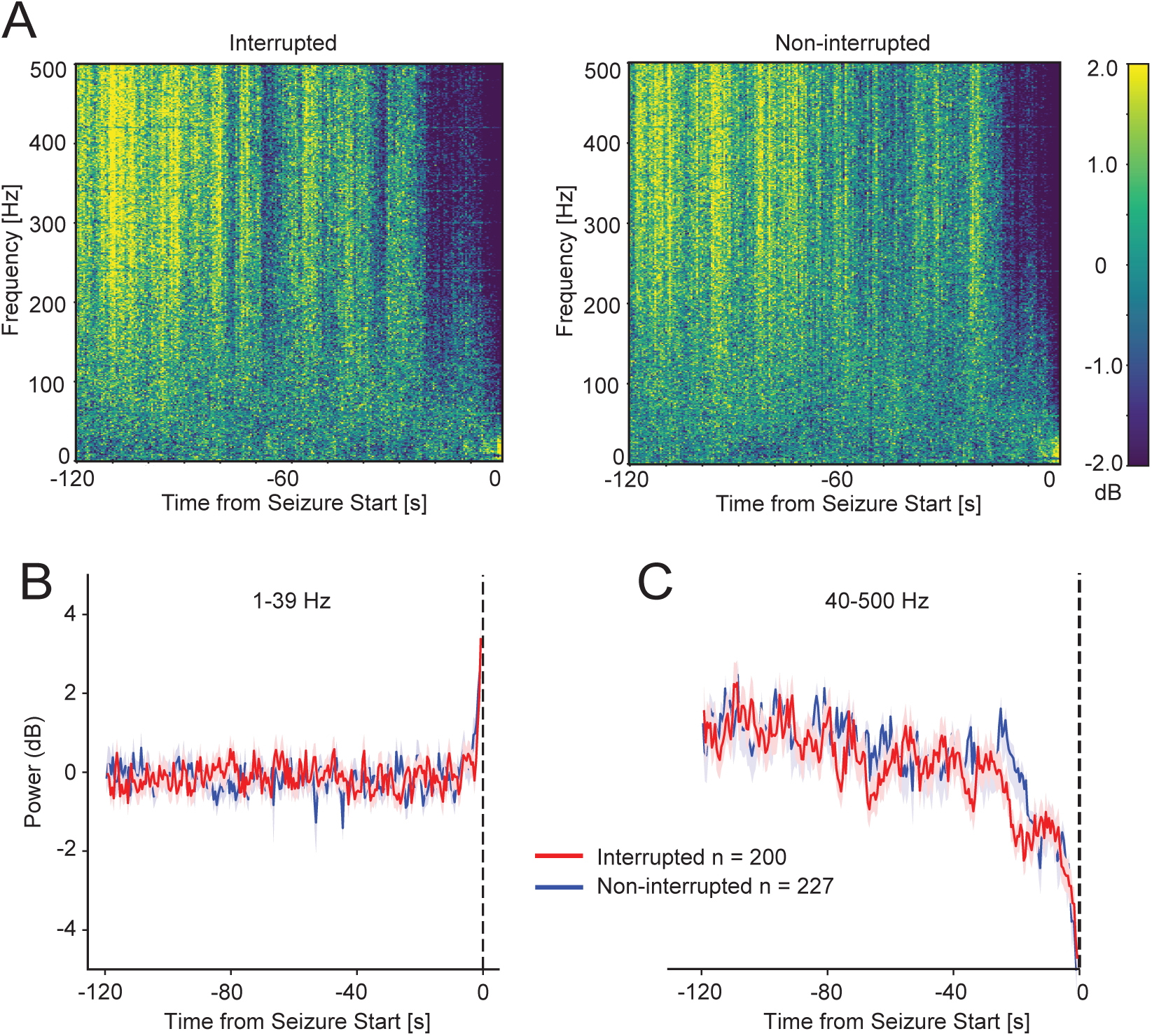
Brain state before seizure was not detectably different between interrupted and non-interrupted seizures. A: Spectrograms of interrupted and non-interrupted seizures calculated over the 120 s preceding seizure start did not demonstrate any obvious differences. B: Power in lower frequency ranges (1-39 Hz) in the 120 s preceding seizure showing similar trajectories for interrupted and non-interrupted seizures. C: As in B but for higher frequencies (40-500 Hz).

## Discussion

In this study we have shown that auditory stimuli associated with a sucrose-water reward can cause early termination (interruption) of spike-wave discharges in the GAERS model. This interruption occurs in approximately 50% of seizures that receive stimulation and is followed by rapid electrographic and behavioral recovery to the non-seizure state. Stimuli that fall earlier in a seizure are more likely to interrupt it, and increased conditioning of stimuli also makes interruption more likely. Electrographic properties differed between interrupted and non-interrupted seizures, both globally and immediately (250 ms) before stimulus delivery. We did not observe electrographic or behavioral differences in the two minutes preceding interrupted and non-interrupted seizures.

### Absence seizure interruption by sensory stimulation

The ability of auditory stimuli to interrupt SWDs is compatible with previous evidence showing that cortical sensory processing persists an absence seizure (Chipaux et al., 2013; Polack, 2016). It matches observations in people with absence seizures (Rajna and Lona, 1989). The variable nature of these interruptions is also in line with previous behavioral observations (Guo et al., 2016). While it has been argued that lessened behavioral severity of rodent SWDs damages their utility as models of human absence seizures (Wiest and Nicolelis, 2003; Taylor et al., 2017; Taylor et al., 2022), both the interruptions and their variable behavioral severity were in fact first demonstrated in people (Mirsky and Vanburen, 1965; Rajna and Lona, 1989; Guo et al., 2016), and we therefore propose that their existence in GAERS provides a means of further investigating the nature and mechanisms of these clinically-relevant aspects of the condition.

### Factors influencing seizure interruption

The factors influencing whether a seizure is interrupted are also of interest. An individual spike-wave discharge is not consistent from beginning to end, in humans or rodents, but rather waxes and wanes in amplitude (Panayiotopoulos, 1999; Pitkänen et al., 2017). Such electrographic waxing often occurs at the onset of an absence seizure (Pinault et al., 2001; Bai et al., 2010); that is, the initial spike-wave complexes of a discharge are less morphologically distinct and lower amplitude than those that follow. In rat (Meeren et al., 2002; Polack et al., 2007) and in human absence seizures (Holmes et al., 2004; Tenney et al., 2013) this may reflect a focal initiation of the seizure that gradually spreads to other brain regions. The fact that earlier stimulus delivery was more likely to precede interruption of seizures both in GAERS and humans (Rajna and Lona, 1989) is compatible with evidence that this lower-amplitude oscillation reflects a less extreme seizure state (Carney et al., 2010; Kumar et al., 2023), that is more easily disrupted and diverted back to a non-seizure brain state. This has implications for potential therapeutic interventions aimed at seizure interruption at discrete time points, for example via closed-loop electrical or (in the future) opto/chemogenetic stimulation (Fan et al., 2017; Wang and Wang, 2017). It is somewhat incongruent with observations that early stimulation is associated with reduced responsiveness in seizures in children with CAE (Sadleir et al., 2008; Guo et al., 2016), but in this case we measure seizure interruption rather than responses during an ongoing seizure.

The observation of an increased proportion of interrupted seizures with increased behavioral training is also promising. It may be that preserved cortical processing during seizures allows an evaluation of the salience of incoming stimuli, akin to a gating process, albeit altered from normal sensory processing (Chipaux et al., 2013; Williams et al., 2016). Depending on salience and intensity, stimuli may be able to provoke a perception, a response, or interrupt an ongoing seizure entirely (Rajna and Lona, 1989; Guo et al., 2016). Again, this has therapeutic relevance – either via alternative neuronal targets for stimulation in sensory pathways (Theyel et al., 2010), or via sensory stimulation for seizure interruption that could select the most salient or “penetrative” stimulus possible in a personalized fashion for each patient.

Electrographic differences between interrupted and non-interrupted seizures may also reflect the overall stability of the ictal brain state. Increased signal power seen in the interruption-resistant seizures overall, as well as in the 250 ms preceding stimulus onset, suggests the basic oscillation of the SWD is more pronounced when interruption-resistance is present. This is perhaps due to a larger number of neurons being recruited into the oscillation and expressing rhythmic activity (Blumenfeld, 2005b; McCafferty et al., 2023) or a greater degree of synchrony between participating neurons.

### Pre-seizure state and interruption

Our findings (or lack thereof) on the relationship between pre-seizure state and likelihood of interruption are far from conclusive as the metrics available to us (EEG spectral properties; lick rate) represent only a partial picture of arousal state. Given the now well-established presence of pre-seizure brain dynamics tens of seconds before the electrographic emergence of a SWD (Pinault et al., 2001; Bai et al., 2010; Carney et al., 2010; McCafferty et al., 2023; Khan et al., 2024), the role of pre-event brain state in determining the nature of sleep spindles (an oscillation with some similarities to SWDs) (Bartho et al., 2014), it is possible that the sensitivity of a seizure to interruption will be at least partially pre-determined. It should be noted however that our negative result matches a study showing that pre-seizure arousal state (awake, drowsy, asleep) had no relationship with responsiveness in children with absence seizures (Sadleir et al., 2008).

### Unanswered questions and future directions

Although the phenomena described here mirror prior observations in humans it would still be valuable to establish their existence in other lab models of absence seizures (Pitkänen et al., 2017) to improve the likelihood of translational relevance. Similarly, an update of Rajna & Lona’s 1989 study on sensory interruption of human absence seizures is long overdue. Our study focuses specifically on the early termination, or interruption, of seizures by sensory stimulation and so the question of consciousness during seizure (Groulx-Boivin et al., 2024) is beyond the scope of discussion here. Interrupting seizures and restoring consciousness during seizure may form separate prongs of a therapeutic attack on the negative impacts of absence seizures. The two are however connected in that sensory interruption requires some degree of neural responsiveness during the seizure – the incoming stimulus must be allowed to impact the cortico-thalamocortical system such that it can shunt it out of the seizure state. We may therefore suggest a human study holistically measuring the relationship between sensation and seizure (Studer et al., 2019; Stenroos et al., 2024), characterizing comprehensively both the ability of a stimulus to interrupt and the perception of that stimulus within and without a seizure.

A comparison of the interrupting qualities of other sensory stimulation paradigms and modalities (e.g. somatic sensation, light) as well as the two candidate factors identified here for influencing interruptions – salience and intensity – may point both to the underlying mechanism of sensory processing and interruption during seizure and to the therapeutic applications most likely to succeed. Finally, the mechanism and pathways dictating the results we have observed here should be investigated in GAERS and other animal models, to answer questions like: how do different neurons along the sensory pathways react to ictal and inter-ictal stimuli? How far does an auditory stimulus get during a seizure – is it processed as normal in the inferior colliculus, the medial geniculate nucleus, the primary auditory cortex, higher order cortex? How does the variable state of cortical and thalamic neurons during a spike-wave complex influence the impact of an incoming sensory stimulus?

Broadly, these results add weight to the increasing corpus of evidence demonstrating variability in the impact of absence seizures in humans and animal models. While this variability adds a new challenge to our study of these seizures it also provides an excellent opportunity to characterize what makes a seizure more or less impactful on the ability of a person with absence epilepsy to navigate and interact with their world, such that therapies and interventions can aim to mitigate or remove that impact.

## Acknowledgements

This work was supported by NIH/NINDS R37 NS100901, R01 NS066974, the Mark Loughridge and Michele Williams Foundation, and the Betsy and Jonathan Blattmachr family.

